# Force localization modes in dynamic epithelial colonies

**DOI:** 10.1101/336164

**Authors:** Erik N. Schaumann, Michael F. Staddon, Margaret L. Gardel, Shiladitya Banerjee

## Abstract

Collective cell behaviors, including tissue remodeling, morphogenesis and cancer metastasis rely on dynamics between cells, their neighbors and the extracellular matrix. The lack of quantitative models precludes understanding of how cell-cell and cell-matrix interactions regulate tissue-scale force transmission to guide morphogenic processes. We integrate biophysical measurements on model epithelial tissues and computational modelling to explore how cell-level dynamics alter mechanical stress organization at multicellular scales. We show that traction stress distribution in epithelial colonies can vary widely for identical geometries. For colonies with peripheral localization of traction stresses, we recapitulate previously described mechanical behavior of cohesive tissues with a continuum model. By contrast, highly motile cells within colonies produce traction stresses that fluctuate in space and time. To predict the traction force dynamics, we introduce an Active Adherent Vertex Model (AAVM) for epithelial monolayers. AAVM predicts that increased cellular motility and reduced intercellular mechanical coupling localize traction stresses in the colony interior, in agreement with our experimental data. Furthermore, the model captures a wide spectrum of localized stress production modes that arise from individual cell activities including cell division, rotation, and polarized migration. This approach provides a robust quantitative framework to study how cell-scale dynamics influence force transmission in epithelial tissues.

## INTRODUCTION

Mechanical interactions at cell-cell interfaces and between cells and the extracellular matrix (ECM) play pivotal roles in tissue organization (Zallen, 2007), developmental morphogenesis (Varner and Nelson, 2014), wound repair (Brugués et al., 2014) and cancer invasion (Friedl and Gilmour, 2009). These collective cell behaviors rely on long-range transmission of mechanical forces, tightly coordinated by mechanical cell-cell coupling and biochemical signaling (Ladoux and Mège, 2017). The maintenance of the balance between cell-cell and cell-ECM mechanical forces are essential for tissue cohesiveness and for cooperative cell movement in swirls, packs, and clusters during many developmental processes. However, little is known about the relative roles of cell motility and intercellular mechanical coupling in coordinating mechanical stress transmission and relaxation across tissue scales.

Forces generated by molecular motors are transmitted by the actin cytoskeleton and to neighboring cells via cadherin-based adhesions or to the ECM via integrin-based focal adhesions. The predominant mechanism of cell force generation occurs via the activity of myosin II on actin filaments, collectively known as the actomyosin cytoskeleton, to generate contractile forces. The actomyosin cytoskeleton, in turn, applies active contractile forces at the sites of integrin-based cell-ECM adhesions. Several studies have implicated that the mechanical crosstalk between actomyosin contractility, cell-cell and cell-ECM adhesions (de Rooij et al., 2005; Maruthamuthu et al., 2011; McCain et al., 2012; Tsai and Kam, 2009) can regulate cell traction forces, cell shape changes, and cellular motile behavior. We and others have shown how epithelial colonies with strong cell-cell coupling can propagate traction stresses across multicellular length scales (Mertz et al., 2012; Mertz et al., 2013; Tambe et al., 2011). In the absence of strong intercellular adhesions, cell traction stresses are primarily localized to cell-cell interfaces (Mertz et al., 2013). While cadherin-based cell-cell forces have been shown to modify traction stress organization and contractility of static epithelial monolayers, we lack a quantitative predictive model for how intercellular tension is propagated to the ECM in highly motile cell colonies with diverse geometries. Progress is limited by the lack of a robust model system that allows precise determination of mechanical forces under controlled environmental conditions, which can be used to build quantitatively accurate cross-scale physical models. Here we address this challenge using an *in vitro* experimental system together with a computational model that enables multi-scale analysis of epithelial mechanics with precise control over cell density, cellular adhesive and motile properties, ECM mechanics and geometry.

Using a combination of Traction Force Microscopy (TFM) (Sabass et al., 2008) and micropatterning on Madin-Darby canine kidney (MDCK) cell colonies, we show that cellular traction forces can vary widely in magnitude and spatial organization for the same geometry and composition of the colony. For MDCK colonies with peripheral localization of traction stresses, we recapitulate results previously reported for single adherent cells (Oakes et al., 2014) and strongly adherent cell colonies (Mertz et al., 2012). In this case, the colony behaves like a macroscopic contractile medium, and its overall mechanical output can be accurately described using a previously developed continuum model (Banerjee and Marchetti, 2012). However, this model is inadequate to describe traction stress organization in colonies with highly motile cells, where cell traction forces are distributed throughout the colony interior. To this end, we develop a dynamic vertex model for motile epithelial cell colonies adherent to a soft elastic substrate. In contrast to purely mechanical vertex models (Farhadifar et al., 2007), our approach accounts for the changes in cell shape and adhesion that occur during cell motion. In quantitative agreement with experiments, our cell-based model predicts a robust relationship between individual cell motility and traction stress localization in large colonies. Furthermore, our model successfully predicts how colony traction patterns can be modulated by internal cell events such as division, polarized motility and collective rotations. Thus, we propose a robust quantitative framework for multiscale analysis that allows us to predict the regulation of mechanical stress transmission from single-cell to tissue level.

## RESULTS

### Colonies with same geometry vary widely in the spatial organization of traction stresses

We used Madin-Darby canine kidney (MDCK) cells as a model system for tissue mechanics. Seeding MDCK cells on collagen-coated TFM substrates yields isolated colonies of ∼10-100 cells. While colonies tend to be circular, they can achieve a variety of shapes due to spontaneous motions and stresses within the colony. Previously, we recognized the roles of geometry in controlling traction stresses in isolated cells (Oakes et al., 2014). In order to control colony shape, we employed identical micropatterning techniques for collagen deposition on TFM substrates. As MDCK cells do not adhere to polyacrylamide (PAA) gels, this constrained cell attachment only to areas defined by the collagen micropatterns.

Surveying a large number of model colonies, we observed that colonies tended to distribute traction stresses in one of two ways. In some cases, traction stress peaks that are confined to the colony periphery and are oriented inward and perpendicular to the colony edge (Figure 1A, left and middle). The second mode of distribution has significant stress peaks in the colony interiors as well as on the periphery (Figure 1B, left and middle). Traction vectors associated with this pattern also display a greater variation in their orientation relative to the colony border (Figure 1B, middle; Movie S1). In both cases the vector sum of all forces throughout the colony is small, less than 10% of the total strain energy, indicating that the traction forces are balanced across the cell island.

**Figure 1:**
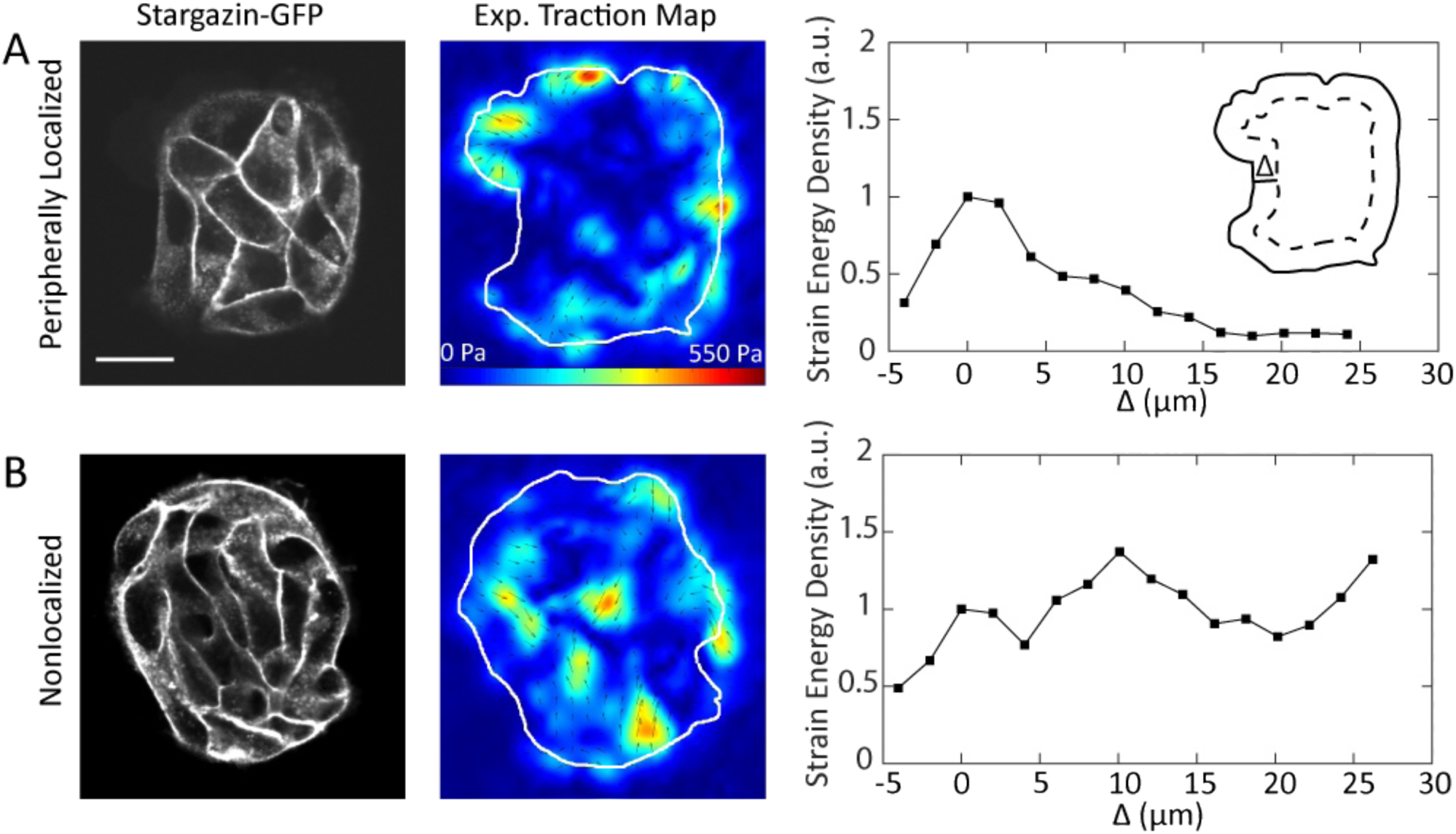
The traction stress distribution in epithelial colonies can vary widely for the same geometry. (A) A colony with traction stresses localized at the colony periphery. (Left) The shapes of cells and the overall pattern shape visualized with the membrane marker Stargazin-GFP. (Middle) Traction stress heatmap for the colony at left – the colony shape is outlined in white. (Right) Strain energy density as a function of Δ, the distance from the colony border, normalized to the value at Δ = 0. The peak at or near Δ = 0 is the defining feature of colonies with peripheral stress localization. (Right, inset) Schematic of the procedure used to generate strain energy density profiles. (B) A colony with a higher proportion of internal traction stresses. (Left-right) Cell and colony shapes visualized with Stargazin-GFP, traction heatmap showing stresses localized to hot spots distributed throughout the colony interior, strain energy density profile that increases with Δ.

To quantify the extent of peripheral localization of stresses, we employed the same procedure used in (Mertz et al., 2012) to plot strain energy density as a function of distance from the colony edge. We divided each colony into a series of concentric contours with width x, which followed the shape of the colony outline such that the ring associated with distance Δ covers every point with a distance between Δ and Δ + x. Because the TFM analysis yields traction footprints with widths on the order of several microns, we included negative Δ values to capture stresses that are exerted outside of the border. We calculated the strain energy of each region by taking the dot product of the traction force with the displacement, and obtained the strain energy density by dividing this quantity by the area of the region. To better compare results from several colonies, we normalized each strain energy density to its value at Δ = 0. The resulting plots provide a quantitative profile of traction stress localization relative to the colony edge. Peripherally localized colonies have a maximum value at or near Δ = 0, with the strain energy density rapidly decaying further into the interior (Figure 1A, right). For colonies with sizeable tractions throughout the interior, strain energy density at Δ = 0 is not necessarily the maximum, and the profile does not decay near the center of the colony (Figure 1B, right).

### Mechanical output of colonies with peripheral localization of traction stresses are identical to single cells

MDCK colonies with peripherally localized traction stresses bore striking resemblance with previously reported traction stress patterns for adherent single cells (Oakes et al 2014) and strongly cohesive keratinocyte colonies (Mertz et al., 2012) that are well described by a continuum model (Banerjee and Marchetti, 2012; Edwards and Schwarz, 2011). Briefly, the continuum model describes the cell colony as an isotropic and homogeneous elastic medium, subject to a uniform contractile stress (force per unit area) due to actomyosin contractility (Materials and Methods). The contractile medium is coupled to a soft elastic substrate via stiff springs that represent adhesion bonds (Figure 2A). This isotropic homogeneous model was sufficient to capture traction stress localization in circularly shaped cells and colonies, where traction stresses spread out evenly along the periphery (Mertz et al., 2012; Oakes et al., 2014). When individual cells are constrained to non-circular adhesive geometries, traction stresses are constrained to regions of high curvature, and an edge contractility parameter was needed for the model to capture the experimental results (Oakes et al., 2014). The extent to which similar traction stress localization would also occur in non-circular model epithelial colonies remained unknown.

**Figure 2:**
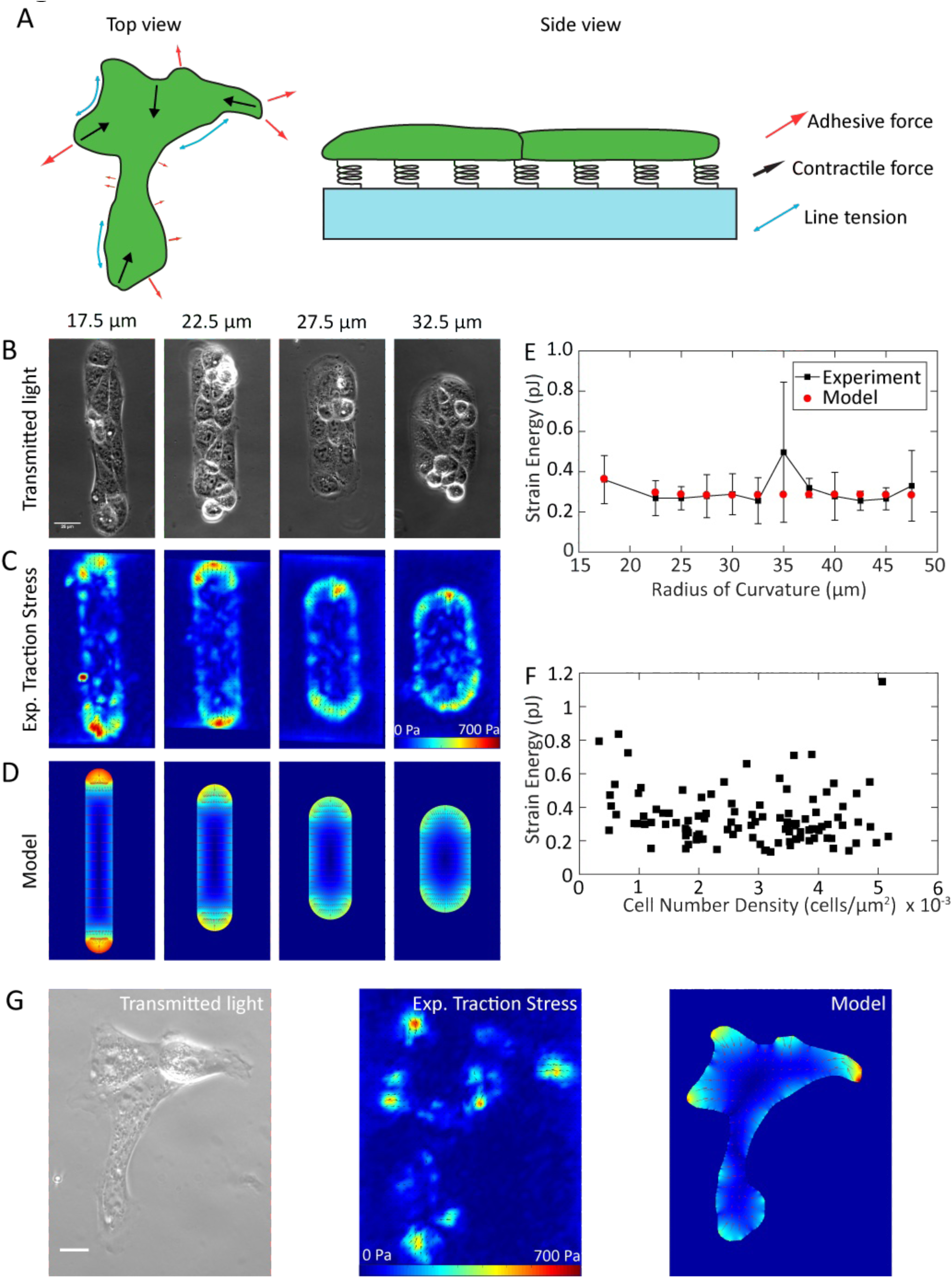
For colonies with peripheral localization of traction stress, a continuum mechanical model quantitatively captures traction stress distribution. (A) Schematic of the continuum mechanical model. (B-D) Peripheral localization of traction stresses in ZO-1/2 dKD MDCK colonies is quantitatively captured by a homogeneous continuum model for cohesive cell colonies. (B) Phase contrast images of ZO-1/2 dKD MDCK cells in stadium-shaped micropatterns of constant area and varying radii of curvature. (C) Traction stress heatmaps for constant area colonies, averaged over n = 4-9 different colonies. (D) Continuum model results for given colony geometries with model parameters: E_cell_ = 6.8 kPa, *σ*_a_ = 780 Pa, and *f*_m_ = 0.3 nN/µm. (E) Colony strain energy is independent of colony shape. Each data point represents average over n = 4-9 colonies. (F) Strain energy does not depend on the number density of cells within a colony. (G) Traction stress organization in unconstrained MDCK colonies can also be described by the continuum model. Left to right: phase contrast, experimental traction map, and continuum model traction map images for an adherent colony on an unpatterned substrate. Model parameters are the same as in (D).

To test whether predictions of this continuum mechanical model (Materials and Methods) were consistent with cell colonies of arbitrary geometries, we used micropatterning to create 8000 µm^2^ stadium-shaped patterns, with radii of curvature of the ends that span from 17.5 µm to 47.5 µm. For these experiments, we used zonula occludens-1/2 double knockdown (ZO-1/2 dKD) MDCK cells (Choi et al., 2016; Fanning et al., 2012), which display stronger peripheral localization than their wild type counterparts. For each adhesion geometry, we acquired phase-contrast images of the cells (Figure 2B) and measure traction stresses transmitted to the substrate. For each geometry, we measured the traction stresses of multiple colonies and calculated the time-averaged stress field (Figure 2C). This revealed that traction stresses are localized to regions of curvature for the model colonies, with the stress vectors oriented inward and perpendicular to the colony edge, similar to what we observed for single cells (Oakes et al., 2014). Using the continuum model, we parametrized the elastic modulus for the colony (6.8 kPa), the colony edge line tension (0.3 nN/µm) and active contractile stress (780 Pa) required to recapitulate the traction stress distribution (Figure 2D) and strain energy (Figure 2E) of colonies with varied geometry. Consistent with our previous findings for single cells, we found the total mechanical work (strain energy) of the colony is independent of the colony geometry for a constant area (Figure 2E). Further, we established that the mechanical work is independent of the cell number density (Figure 2F). This indicates that the tissue-scale mechanical properties that can be described with the continuum model are determined by the global tissue shape and mechanical properties, and are independent of cell-scale properties such as their shapes and density.

To demonstrate the applicability of this continuum model to epithelial colonies of arbitrary geometries, we tested its utility in predicting force localization of an unconstrained colony. In Figure 2G, phase contrast and traction force measurement of a small colony containing four MDCK cells are shown, chosen for its highly extended shape. Traction stresses are localized at the periphery, and are further concentrated in small, high curvature regions. Using the parameter values benchmarked from the micropattern and TFM experiments, the continuum mechanical model successfully predicts the spatial localization and magnitude of the traction stresses for this colony shape (Figure 2G, right). Thus, colonies with peripheral localization of traction stresses with varying geometries are well characterized by a continuum model that has a bulk contractility and an edge tension, similar to what we observed previously for single cells (Oakes et al., 2014). Taken together, these results suggest a regime in which adherent cell colonies exhibit identical mechanical behavior as single cells, such that its global mechanical output is independent of individual cell properties.

### Active Adherent Vertex Model (AAVM) for traction stress prediction in dynamic epithelial colonies

Our experimental data suggest another regime for mechanical force transmission in colonies, where traction stresses are distributed throughout the colony interior (Figure 1B,D,F). One underlying mechanism of interior stresses arises from reduced cell-cell coupling (Mertz et al., 2013). However, we sought to explore the extent to which cell motion within a confluent tissue could underlie interior traction localization. To predict traction stress localization in dynamic colonies, we developed Active Adherent Vertex Model (AAVM) for predicting mechanical stresses in adherent cell colonies. This approach allows explicit control over the dynamic mechanical properties of individual cells, which are not accessible by a continuum description of colony mechanics. In contrast to purely mechanical cell-resolution Vertex Models (Barton et al., 2017; Bi et al., 2015; Farhadifar et al., 2007; Fletcher et al., 2014; Honda and Eguchi, 1980) or Cellular Potts Models (Albert and Schwarz, 2016; Graner and Glazier, 1992), AAVM explicitly accounts for the coupling between mechanical forces that drive cell motion and the kinetics of cell-matrix adhesion binding/unbinding, active cell motility from cell protrusions, retractions, and cell polarity fluctuations (Figure 3A, Figure S9). We model the cell colony as a confluent two-dimensional monolayer, where the geometry of each cell is described by a polygon (Figure 3A, Materials and Methods). To allow the formation of curved cell shapes, we subdivide the cell contour into a large number of linear segments. Physical interactions between cells are modelled following the vertex model for epithelial tissues, whose mechanical energy arises from cellular elasticity, cortical tension, cell-cell adhesion, and actomyosin contractility (Farhadifar et al., 2007).

**Figure 3:**
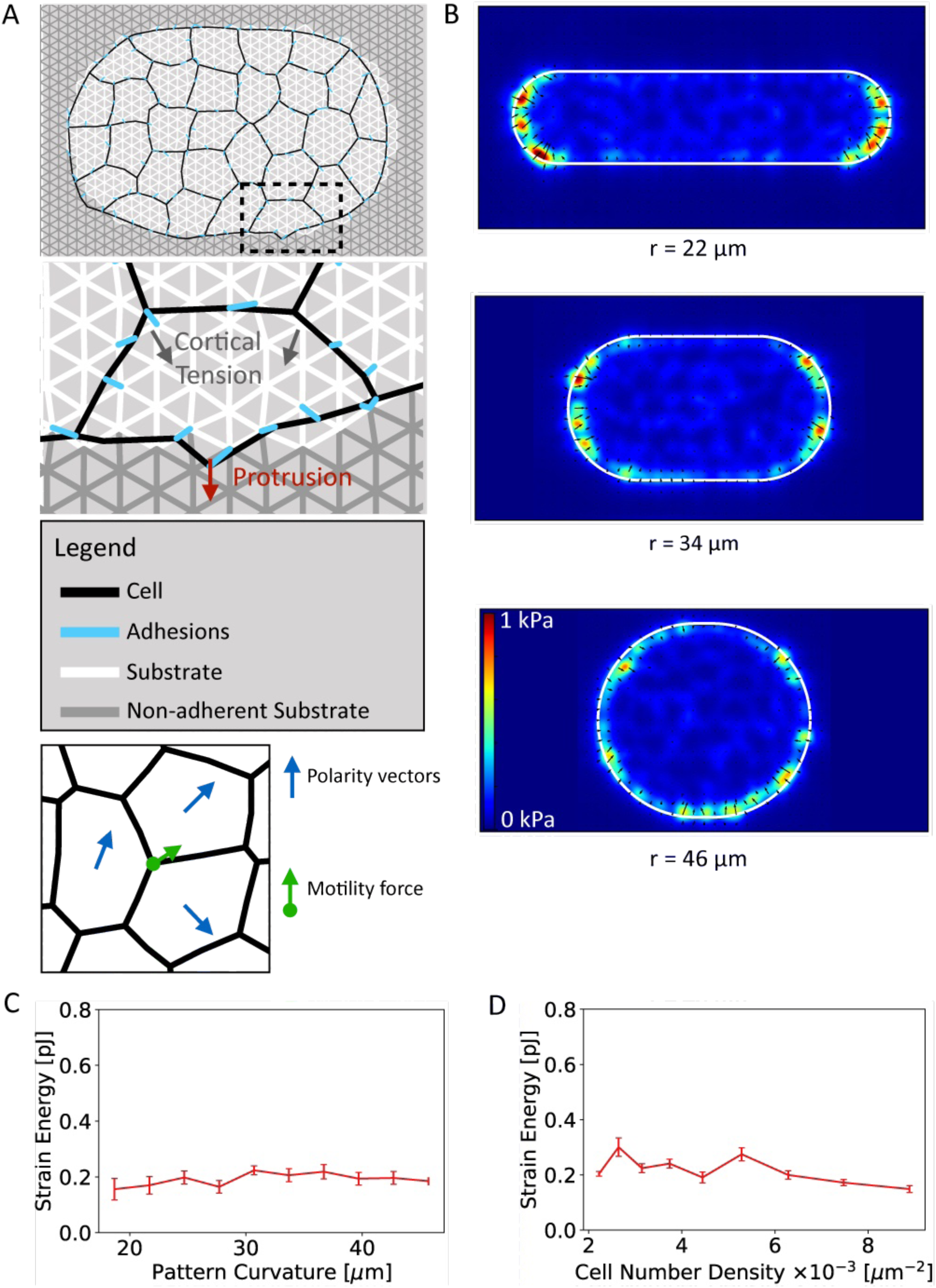
Active Adherent Vertex Model for epithelial cell colonies can be benchmarked to experiments to capture spatial variations in traction stress. (A) Simulation image for a cell colony on a micropattern (top) and a zoomed in region illustrating the mechanical forces acting on adherent cells (bottom). (B) Time averaged traction stress maps for varying curvature radii of the micropattern: r = 22 µm (top), r = 34 µm (middle), and r = 46 µm (bottom). (C) Strain energy as a function of pattern curvature for a fixed area. (D) Strain energy as a function of cell density for fixed pattern shape.

The soft elastic substrate is modelled by a triangular mesh of harmonic springs, which are anchored to cell vertices via stiff springs (Figure 3A), representing focal adhesion complexes. The cell-substrate adhesions bind and unbind stochastically at fixed rates. We model the confining effect of the micropattern by disallowing adhesions outside a pre-defined geometry of the substrate. Cells in the interior of the colony move with a speed *v*_0_ in the direction of its polarity (Materials and Methods). Cells on the boundary of the colony assemble protrusions at their external edges, which push the cell vertices forwards that subsequently bind and pull on the substrate. Each cell vertex evolves in time following an overdamped equation of motion, where the cell vertex velocity is proportional to forces resulting from total mechanical energy of the colony, cell-substrate adhesions and active cellular motility (Materials and Methods).

By benchmarking the physical parameters of AAVM (Supplemental Material) we successfully captured the experimental results for cohesive cell colonies in Figure 3 (Movie S2). To this end, we set the internal motility speed, *v*_0_, of the cells to be comparable to the low motility speed for ZO-1/2 dKD MDCK cells. In quantitative agreement with TFM experiments and the continuum model, we find that traction stresses localize around the curved periphery of the cell colony. Furthermore, as the radius of curvature decreases, the local traction stresses increase in magnitude as they are concentrated in a smaller region (Figure 3B). By measuring the strain energy density as a function of distance from the colony edge, we find that most strain energy is applied around the periphery of the micropattern, which decays with distance from the edge of the colony (Figure 3C, Figure S1). While the magnitude of local traction stresses changes with curvature, the model captures the experimental result that the total strain energy is independent of pattern curvature (Figure 3C). With this same set of parameters, our model accurately predicts the linear dependence of the total strain energy on the colony area (Figure S2), as observed for single cells and cohesive colonies (Mertz et al., 2012; Oakes et al., 2014). By altering the preferred cell area for a fixed area of the micropattern, we can control the number density of cells in a given colony. As a result, we found that the colony strain energy is independent of cell number density (Figure 3D), in quantitative agreement with our experimental data (Figure 2).

### Enhanced cell motility promotes traction stress localization in the colony interior

Since AAVM explicitly accounts for individual cell properties, including tension, adhesion and motility, we sought to investigate how the interplay between active cell motility and intercellular mechanical coupling regulates stress transmission in the colony. Motile behavior of cells can be induced by increasing the magnitude of the self-propulsion speed, *v*_0_, or by inducing fluid behavior of the colony. Fluidity can be induced by lowering the effective mechanical tension at cell interfaces, by either decreasing cortical tension or by increasing cell-cell adhesion energy per unit length. It has been recently shown that increase in cell shape anisotropy can reduce the effective tension at cell interfaces, leading to fluidization of a jammed tissue (Bi et al., 2015). Cell shape anisotropy can be characterized by the dimensionless preferred shape index, 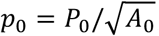, where *P*_0_ is the preferred cell perimeter and *A*_0_ is the preferred cell area (Materials and Methods). While the molecular mechanisms regulating spontaneous changes in *p*_0_ remain to be characterized, here we treat *p*_0_ as a control parameter that tunes tissue fluidity by promoting local cell movement and mechanical stress relaxation by cellular neighbor exchanges. Tissue fluidity can be further increased by increasing cell self-propulsion velocity, which increases the propensity of cell-cell rearrangements (Bi et al., 2016).

As we increase cell motility (by increasing *v*_0_) and tissue fluidity (by increasing *p*_0_) we observe that higher traction stresses are generated in the colony interior (Figure 4A; Movie S3). This leads to delocalization of strain energy away from the colony periphery (Figure 4B), and appears in correlation with enhanced cell movement (Figure 4C). To quantify the dependence of traction stress distribution on cell motility, we systematically varied the self-propulsion speed, *v*_0_, and the cell shape index, *p*_0_. Over all simulations, we find that the decay length λ of traction stresses from the periphery in relation to cell motility increases monotonically as the mean instantaneous velocity of the cells increases (Figure 4D). This quantity is obtained by fitting an exponential function to strain energy density profiles obtained from the earlier annular analysis: *SE = Ae*^*–Δ/λ*^ (Materials and Methods). The resulting decay constant provides a length scale over which the strain energy decreases with distance from the colony border, such that a higher value corresponds to a higher proportion of interior stresses. Increasing either the self-propulsion speed or the cell shape index also leads to an increase in the frequency of cellular neighbor exchanges or intercalation rates (Figure 4E, Figure S3). This increased rate of cell intercalations exhibits strong positive correlation with higher internal motility and lower strain energy localization at the colony periphery.

**Figure 4:**
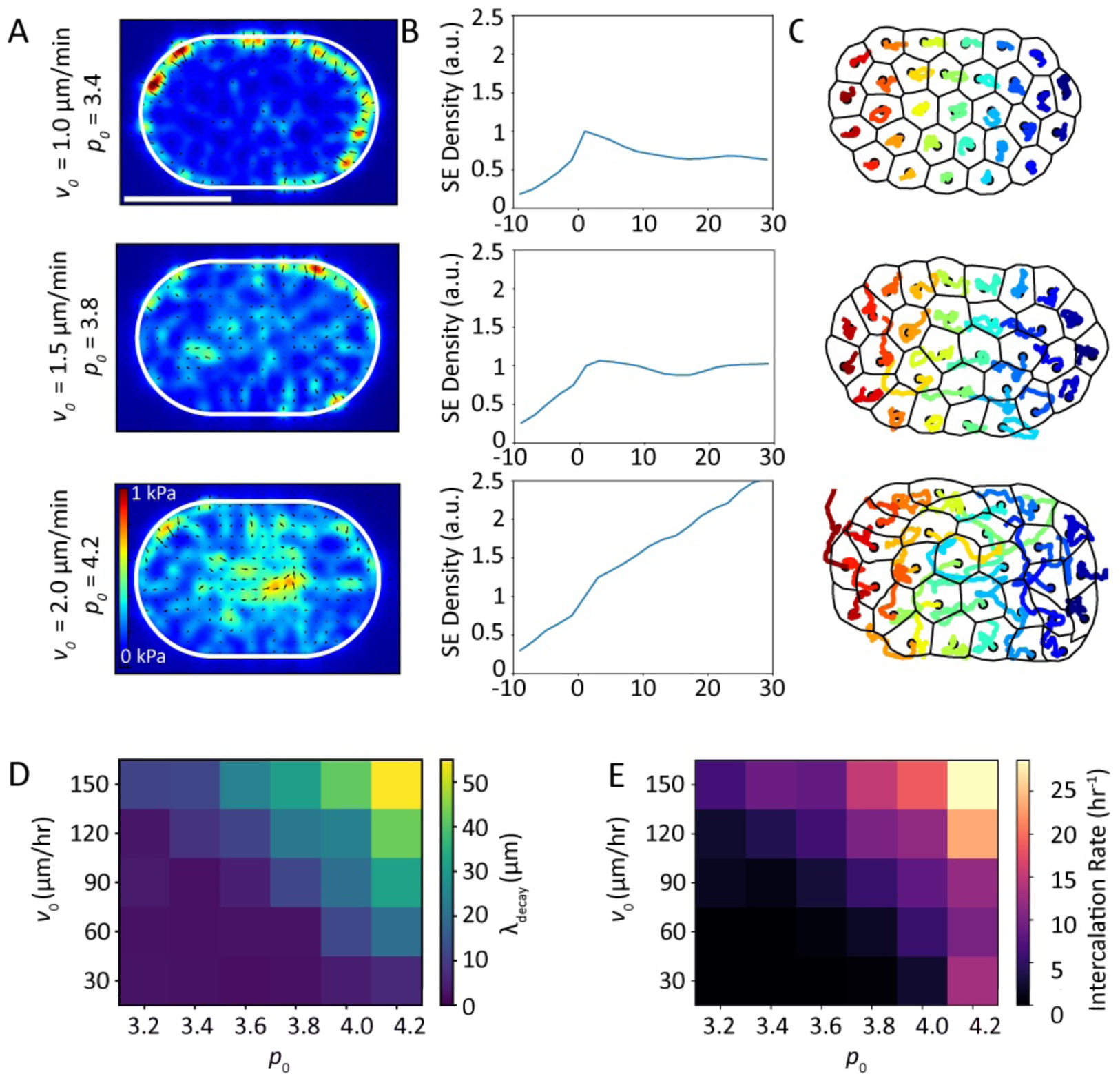
Increased cell motility promotes strain energy localization throughout the cell colony. (A) Time-averaged traction stress maps, (B) averaged strain energy density normalized by the boundary value, (C) and cell trajectory plots for low motility cells, v_0_ = 60µm/hr, P_0_= 3.4 (top), medium motility cells, v_0_ = 90µm/hr, P_0_= 3.8 (middle), and high motility cells, v_0_ = 120µm/hr, P_0_= 4.2 (bottom). Δ is the distance from the boundary and is defined as negative outside of the cell colony, and positive inside the cell colony. Scale bar represents 50 µm. (D) Traction stress decay length λ and (E) rate of cell neighbor exchanges (intercalations) for varying cell motility speed and cell shape index. The decay length λ is measured by fitting *SE = Ae*^*–Δ/λ*^ to the relevant strain energy density profiles.

Our experimental results confirmed the model predictions regarding the relationship between individual cell motility and peripheral traction stress localization. Working under the hypothesis that more motile, fluidized colonies may feature more spatially disperse traction stresses, we considered two representative micropatterned colonies of comparable geometries, which were selected based on differences in mean cell speed over the course of 3.5 hours (Figure 5A,B and Movies S4,S5). Figure 5A shows representative phase contrast and traction heatmap images from a colony where cells have an average instantaneous velocity of 7.92×10^-2^ µm/min, and travel an average of 16.1 μm over 3.25 hours. Cell movement is indicated by the nuclear tracks overlaid on the phase contrast (Figure 5A, left panel). In this colony, the traction stresses are visibly confined to the colony periphery (Figure 5A, middle panel), an observation that is confirmed by the strain energy profile for that time point (Figure 5A, right panel). The same analysis for a colony with considerably higher cell motility, with instantaneous velocity of 0.197 µm/min and mean path length of 48.3 µm over 3.25 hours (Figure 5B, left panel), yields different results. In this case, traction stress peaks are found on both the colony periphery and interior (Figure 5B, middle panel), and the strain energy profile reveals a slow decay in strain energy density from the colony border (Figure 5B, right panel). These differences in traction stress distribution were concordant with the vertex model predictions (Figure 4A) - as the average cell motility in a colony increases, traction stresses reorganize to the colony interior.

**Figure 5:**
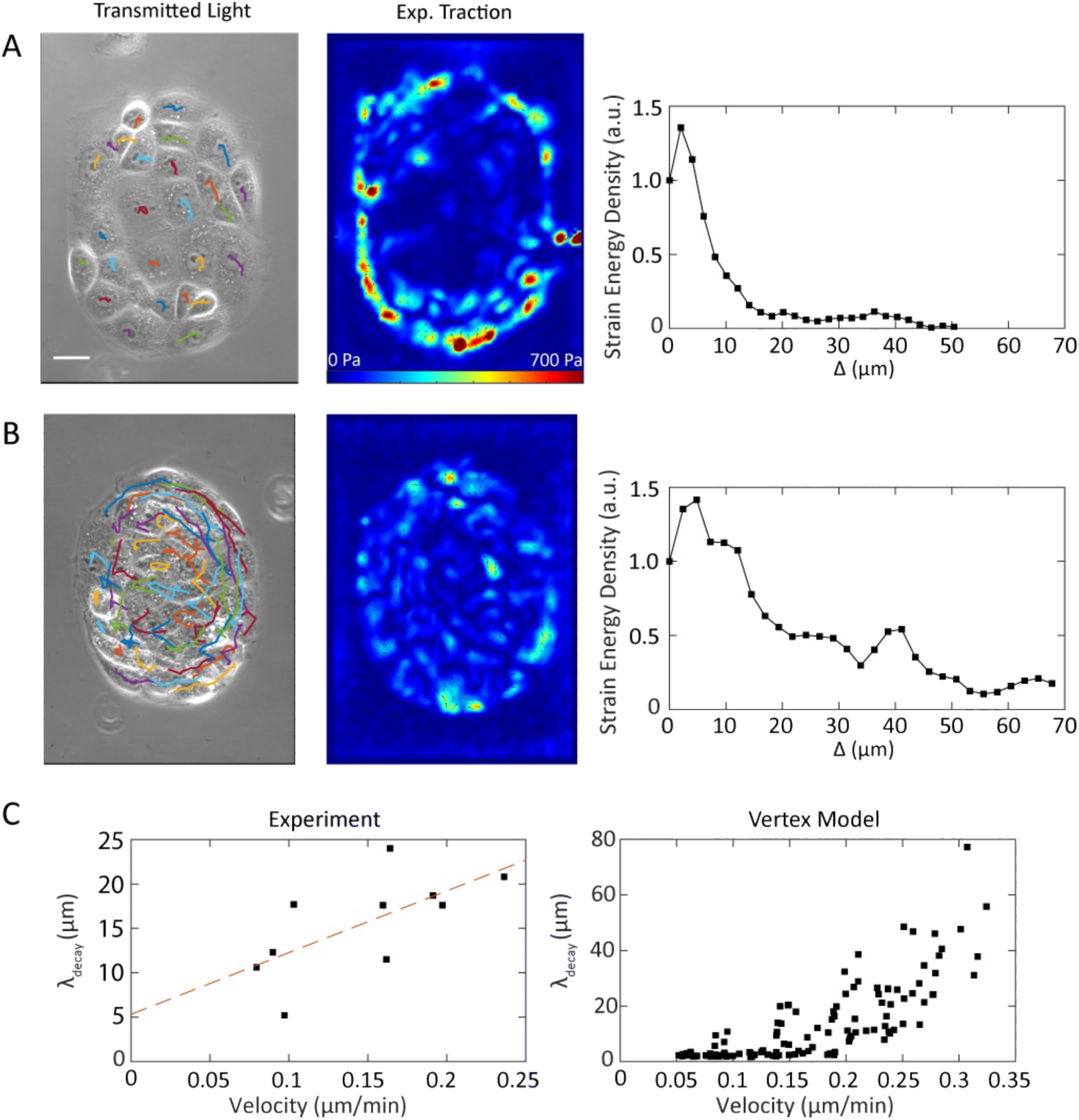
Colonies with interior traction stresses exhibit high degree of individual cell motility. (A,B) Colonies with peripheral localization of traction stresses have cells that appear jammed and do not reorganize traction stresses rapidly. By contrast, colonies with interior traction stresses have highly motile cells and exhibit dynamic peaks in traction stresses. From left to right: Phase contrast images and nuclear tracks over 4 h for two colonies of ZO-1/2 dKD MDCK cells, representative traction maps from each colony, and the strain energy density profiles corresponding to the traction maps. Scale bar = 25 μm. (A) A colony with low motility (mean cell speed = 0.079 µm/min) and traction stresses localized to the colony periphery. (B) A colony with higher motility (mean cell speed = 0.236 µm/min) and a higher proportion of internal stresses. (C) Colonies with higher motility have a longer decay length for strain energy as a function of distance from border. This relationship holds in experiments (left) and is predicted by the vertex model (right) to be robust over a wide range of motilities, using each simulation used in Figure 4D. The red line in the left panel of (C) shows the linear fit of the data shown with slope = 69.53 and y-intercept = 5.313. The radii of curvature for the colonies shown in (A) and (B) are 65 µm and 70 µm, respectively.

We further quantified the relationship between cell motility speed and traction stress organization using the traction stress decay length, λ. We found the average decay length for movies of several colonies, ranging in size from 8000-20000 µm^2^, and found that an increase in cell speed was correlated with a higher decay length (Figure 5C, left panel). We thus experimentally validated the theoretical prediction that traction stress dispersion should accompany enhanced cell motility (Figure 5C, right panel).

### Active cell behaviors coordinate localized stress production modes

Our bottom-up computational modeling and experiments enable us to bridge the gap between single cell dynamics and multicellular mechanical output in adhesive micropatterns. Using this integrative approach, we related the collective modes of traction stress generation in epithelial colonies with the motile and adhesive behaviors of individual cells (Figures 2-5). To further exploit the predictive power of the AAVM in understanding heterogeneous force transmission in epithelial colonies, we examined two frequently occurring active processes: cell division within a monolayer (Figure 6), and spontaneous rotational motion of a colony (Figure 7). In a typical mitosis event occurring within a colony, there is significant lateral constriction that occurs prior to a cell division (Figure 6A). We further observed that traction stresses gradually reorganize to the region around the mitotic cell (Figure 6B, left three panels) and dissipate after cytokinesis (Figure 6B, right two panels). This result, while limited spatially by the finite resolution of the TFM analysis, is concordant with published results for single cells (Tanimoto and Sano, 2012). This provides an active mechanism of stress relaxation and movement that locally fluidizes the colony.

**Figure 6:**
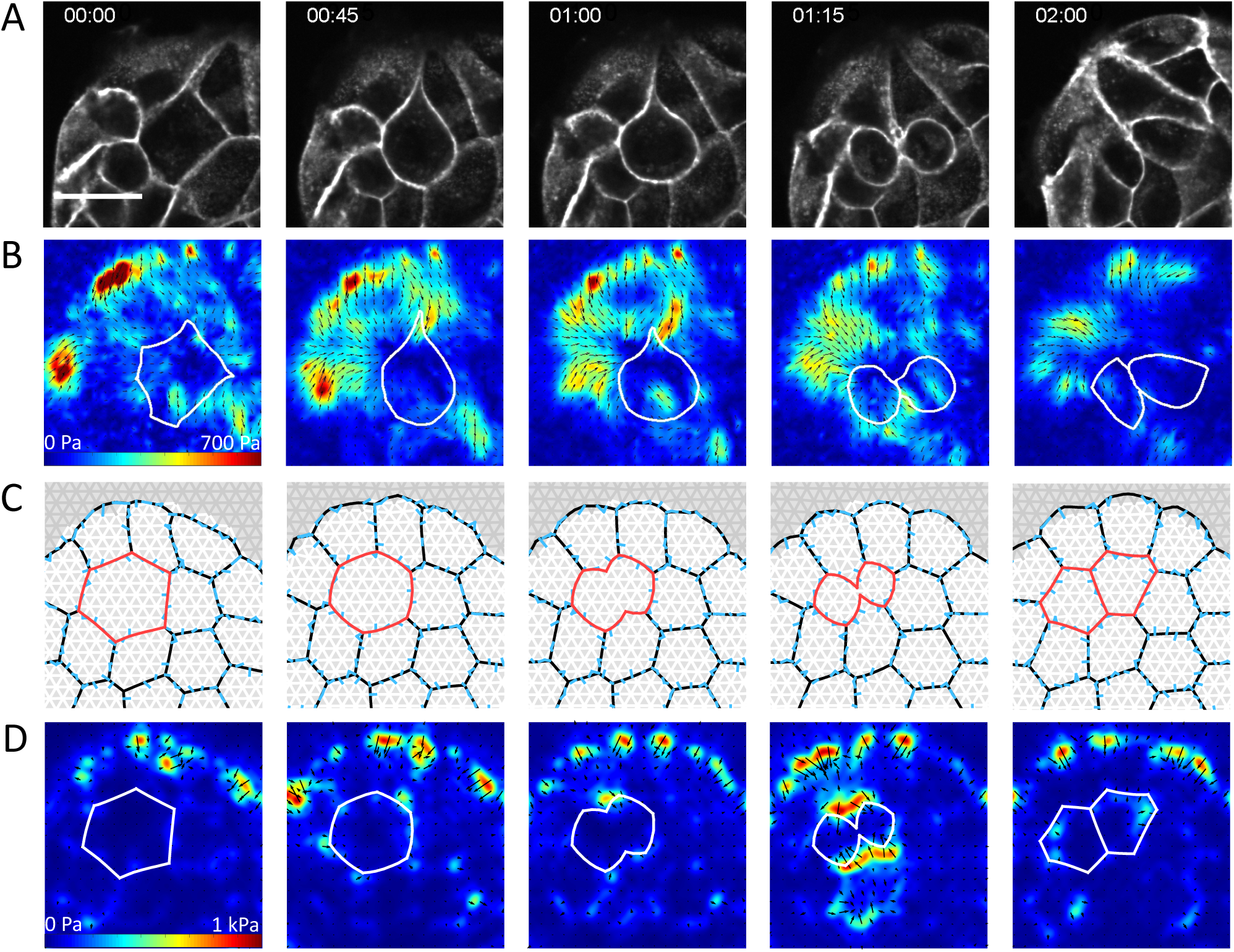
Traction stress localization during cell rounding and division. (A) Stargazin-GFP images showing a cell (outlined in white) contracting and dividing. Scale bar = 10 µm. (B) Traction maps corresponding to (A). As the cell contracts, traction stresses are exerted just exterior to the mitotic cell, directed inward towards the plane of division. Once the cell finishes dividing, the stresses begin to dissipate. (C) Cell shapes during a simulated mitotic event in the vertex model, for a low motility cell colony, *v*_0_ = 30*µm/hr*, *p*_0_ = 3.6. The dividing cell is highlighted in red. Full details of the cell division implementation are given in the Supplementary Material. (D) Traction maps corresponding to (C).

**Figure 7:**
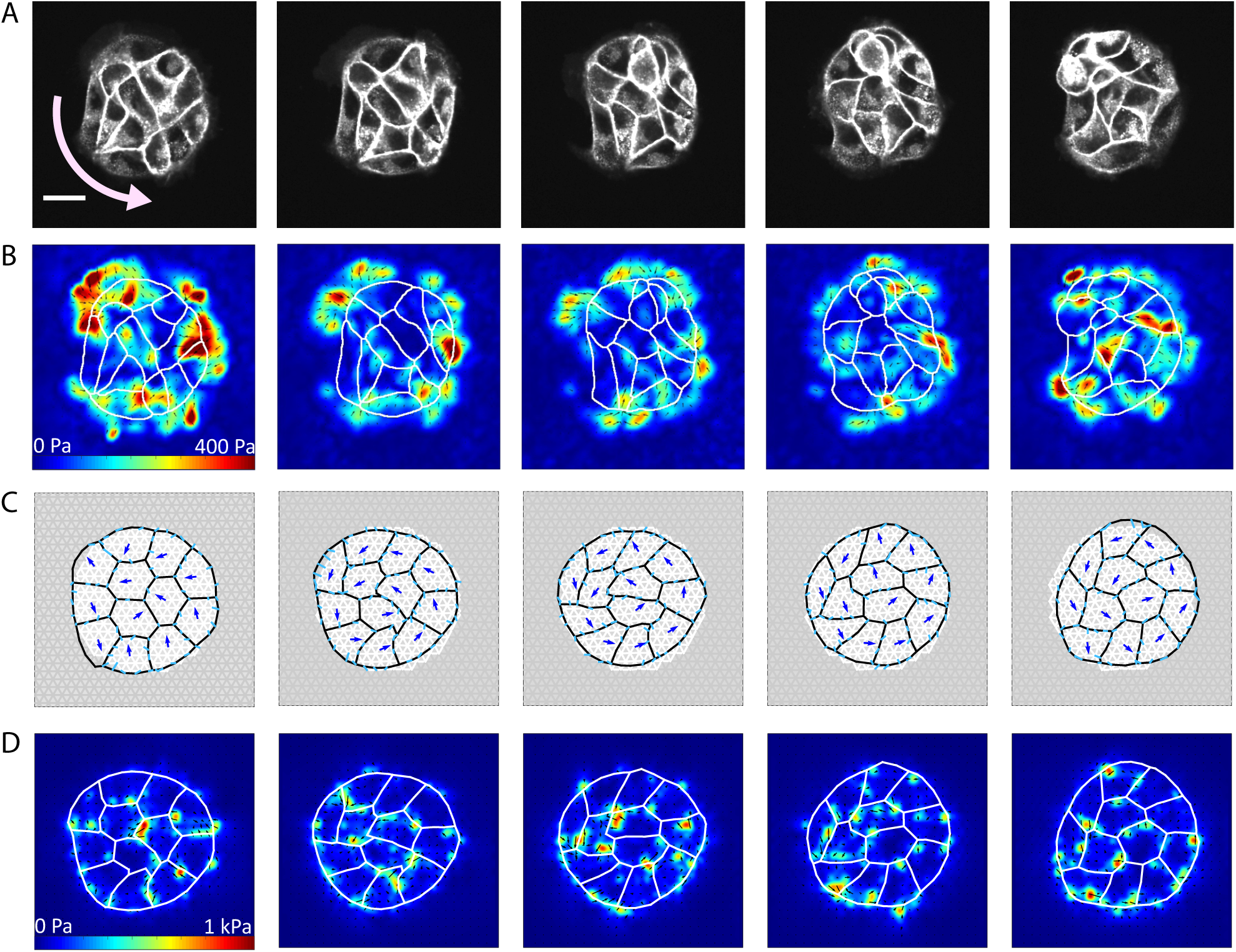
Internal traction stresses form during colony rotation. (A) Stargazin-GFP images of a colony on a circular pattern (radius of curvature = 47.5 µm) rotating counter-clockwise about the pattern center. Scale bar = 25 µm, curved arrow indicates direction of motion. (B) Traction maps corresponding to (A). (C) Cell shapes during rotation simulated by the vertex model, for high motility, v_0_ = 90µm/hr, P_0_= 3.8. Blue arrows show the direction of cell motion. Cell polarities align with cell velocity with timescale τ_v_ = 30 mins, and turn away from the micropattern boundary with timescale τ_b_ = 6 mins. Full details can be found in the Supplementary Material. (D) Traction maps corresponding to (C).

To capture this behavior in AAVM, we implemented the mechanics of cell division in four steps (Figure S4): (i) a *growth phase*, where the preferred area of the dividing cell is doubled (highlighted in red in Figure 6C); (ii) a *cell rounding phase*, where an increased line tension on the cell periphery enable cells to adopt a circular shape (Figure S4B); (iii) a *constriction phase*, where an increased tension at the cell equator results in ingression of the cleavage furrow, around which large traction stresses localize (Figure S4B, Figure 6D); and (iv) *a splitting phase*, where the constricting cell separates into two daughter cells (Figure 6C). Following cell division, the two daughter cells relax their mechanical energies, which results in traction stress dissipation as the cell shapes reach their equilibrium configurations (Figure 6D, Figure S4A). This example illustrates how the active vertex model can be used to capture active stress generation and relaxation due to localized cell shape changes (Movie S6). In particular, local increases in contractility required to change cell shape prior to division are balanced by localized traction stresses generated by neighboring cells.

Next, we examined dynamic organization of traction stresses in colonies undergoing spontaneous rotational motions. The ability of epithelial cells to undergo collective rotations has been proposed to play an important role in acini formation, tissue polarity, and embryogenesis (Tanner et al., 2012). As reported previously (Deforet et al., 2014; Doxzen et al., 2013; Notbohm et al., 2016), small colonies of epithelial cells can often show spontaneous collective rotational motion, with few internal rearrangements (Figure 7A). The limited number of internal rearrangements corresponds to high cohesiveness (low *p*_0_), which is essential for efficient transmission of contact guidance cues from peripheral to interior cells. Based on our findings in Figure 4E, lower fluidity (low *p*_0_) would promote a higher degree of peripheral stress localization. Instead, our traction force measurements show abundant internal traction stresses (Figure 7B).

To recapitulate coordinated cell motion, we simulated a cohesive cell colony with low internal motility (v_0_ = μmhr^-1^,p_0_ = 3.6) where the cell polarity vectors now align with cell center velocities and are repelled by the pattern boundary (see Supplemental Material, Movie S7). The interplay between polarity alignment with local motion and confinement within the pattern results in a coordinated spontaneous rotational motion of the colony (Figure 7C). In the simulations, we find large internal traction stresses localize as the colony coherently rotates (Figure 7D). These localized traction forces accumulate at cell-cell junctions, where they are balanced by traction forces in neighboring cell junctions (Figure S5). These stresses are subsequently dissipated as the cell polarity vectors change in direction during synchronized rotation. This underscores that internal traction stresses can result from coordinated movement due to long-range transmission of mechanical cues by adhesive epithelial cells.

## DISCUSSION

Our work established the relative roles of cell motility and cell-cell interactions on force transmission in multicellular colonies. Through integration of experiments, bottom-up and top-down modeling approaches, we have shown that colony geometry, contractility, cell shape changes, and active motion of constituent cells can profoundly alter the spatial organization of mechanical stresses transmitted to the extracellular matrix. These results provide a robust framework to characterize the role of cell density, motility, geometrical and mechanical cues on collective cell behavior in adhesive environments.

Our results indicate that epithelial colonies organize traction stresses either peripherally or internally (Figure 1) depending on the motile and adhesive behaviors of individual cells. First, we consider the case of cohesive cell colonies exhibiting strong peripheral localization of traction stresses. Consistent with previous results in keratinocytes (Mertz et al., 2012), we find that internal stresses counterbalance in strongly coherent and immobile colonies of cells such that traction stresses become localized to the periphery (Figure 2). The resultant mechanical work performed by the colony on the substrate is independent of the colony shape or cell density, but is proportional to changes in the colony spread area. These findings are consistent with the physical model of a colony described by a characteristic contractility (Figure 2A,D).

Continuum models that treat the multicellular colony as a uniform contractile medium (Mertz et al., 2012) capture this peripheral localization for circular shaped colonies. In previous work on single adherent cells (Oakes et al., 2014), we found that an additional edge contractility was required to capture the traction localization to regions of high curvatures for cells of arbitrary shapes. Here, we find that peripheral contractility is also needed to describe the distribution of traction stresses in multicellular colonies of arbitrary shapes. This edge contractility, characterized by a line tension, naturally arises due to the absence of cell-cell adhesion at free edges, resulting in traction stress localization to regions of curvature. However, this model was insufficient to describe the mechanics of cohesive colonies with sizeable tractions in the interior that fluctuated in space and time (Figure 1B, 5B). This necessitated a cell-based model, AAVM, that accounts for individual cell activity, motility, shape and mechanics (Figure 3A).

While previous work showed that traction stress localization within the colony interior could be explained by loss of cell cohesion (Mertz et al., 2013), here instead the cells remained cohesive over time, exhibiting local dynamic behaviors. We observed that cells within colonies tend to move, exchange neighbors, and change their individual shapes. We therefore developed a new active adherent vertex model that allowed us to capture the dynamic properties of these adherent colonies. By coupling the AAVM to an elastic substrate, we predicted that internal cell motility leading to effective tissue fluidization resulted in a high degree of internal localization of traction stresses (Figure 4). This result was confirmed by our experimental data which showed that the length scale for traction stress penetration increased with increasing cell motility (Figure 5). Our model also captures the traction stress field around a motile cell traversing the colony, which likely arises from weakening of cell-cell adhesions (Figure S6).

Aside from motility driven fluidization, traction stress fluctuations in the bulk of cohesive colonies also occurred due to cell division and rotations. Contractility driven shape changes in dividing cells can locally accumulate traction stresses. These stresses are subsequently dissipated following daughter cell separation, once cell-cell cohesion is established (Figure 6). We also found traction stress dispersion in colonies with low internal motility but where there was highly correlated rotational motion (Figure 7). These persistent rotations have been previously observed in small cohesive colonies in adhesive micropatterns (Deforet et al., 2014; Doxzen et al., 2013; Notbohm et al., 2016). Consistent with previous findings, we find that traction stresses periodically accumulate and dissipate at cell-cell junctions of rotating colonies, arising from coherent transmission of polarity cues by contact guidance. Thus, colonies with the same geometry and adhesive properties can exhibit a rich variety of dynamic properties due to active mechanical behaviors of constituent cells.

In conclusion, we have shown that an interplay between cell motility and cell-cell adhesive interactions can tune the dynamic mechanical properties of cohesive colonies in confined adhesive environments. In doing so, we have developed a quantitative accurate bottom-up model for dynamic epithelial colonies that allows us to predict patterns of collective motion and traction stress generation in diverse conditions. In particular, we show that local active cell behaviors (motility, intercalations, division) in cohesive tissues can induce heterogeneous properties that cannot be captured by simple continuum models. In the absence of active cell dynamics, however, immobile colonies transmit forces like a continuum medium, akin to single adherent cells. These results provide a quantitative framework to predict collective dynamic and mechanical states of epithelial tissues from their emergent patterns of force transmission.

## MATERIALS AND METHODS

### Cell Culture

Madin-Darby Canine Kidney (MDCK) cells were cultured in DMEM media and supplemented with 10% FBS (Mediatech; Corning, Corning, NY), 2 mM L-glutamine (Corning), and penicillin-streptomycin (Corning). To visualize cell shapes, wild type MDCK cells were transfected with plasmid DNA constructs encoding for Stargazin-GFP (courtesy of M. Glotzer, University of Chicago, Chicago, IL). These cells were sorted and found to stably express Stargazin-GFP, with no noticeable loss in marker expression after > 20 passages. ZO-1/2 KD MDCK (dKD) cells were provided courtesy of M. Peifer, University of North Carolina, Chapel Hill, NC.

### Traction Force Substrates

PAA substrates were prepared as previously described (Oakes et al., 2014). Gels with a Young’s modulus of 8.4 kPa were prepared by first making a mixture of 3.125 mL 40% acrylamide (Bio-Rad Laboratories, Hercules, CA), 0.833 mL 2% bis-acrylamide (Bio-Rad), and 1.042 mL water. To this were added 5 µL 110-nm sulfate-modified fluorescent microspheres (Invitrogen, Carlsbad, CA).

### Collagen Micropatterning

Micropatterned substrates were prepared using the same ultraviolet illumination-mediated procedure described in Oakes et al., 2014. Briefly, a chrome-plated quartz photomask (Applied Image Inc., Rochester, NY) was cleaned with water and wiped with 0.3 mL hexane (Sigma-Aldrich, St. Louis, MO) to induce hydrophobicity on the photomask surface. Polyacrylamide gel mixtures were polymerized for 40 minutes between the photomask and prepared glass coverslips with APS and TEMED as radical initiator and co-initiator, respectively. Once the gel was polymerized, the photomask was placed in a UVO-Cleaner 342 (Jelight, Irvine, CA) and illuminated with a combination of 185- and 254-nm ultraviolet light for 90 s. The coverslip and gel were then removed from the photomask by submerging the entire complex in water and gently detaching a corner with a razor blade. Gels were incubated for 10-15 minutes in a solution containing 5 mg/mL EDC (Thermo Fisher Scientific, Hampton, NH) and 10 mg/mL NHS (Thermo Fisher Scientific) kept at acidic pH with MES buffer. The EDC-NHS solution was aspirated and replaced with a solution containing 0.5 mg/mL collagen-I in an MES buffer for 40 minutes. Through this procedure, collagen only crosslinks to regions exposed to UV light as defined by the photomask. Gels were washed 3x for 5 minutes in PBS solution before cells were plated.

### Microscopy and Live Cell Imaging

Cells were imaged on an inverted microscope (Ti-E; Nikon, Melville, NY) with a confocal scanhead (CSU-X; Yokogawa Electric, Musashino, Tokyo, Japan), laser merge module containing 491, 561, and 642 laser lines (Spectral Applied Research, Richmod Hill, Ontario, Canada) and an Andor Zyla sCMOS camera (Belfast, Northern Ireland, UK). METAMORPH acquisition software (Molecular Devices, Eugene, OR) was used to control the microscope hardware. Images were acquired using a 40x 1 NA Plan Apo oil-immersion objective. Samples were mounted on a live imaging chamber (Chamlide, Seoul, Korea) and maintained at 37°C. For live cell imaging, DMEM was supplemented with 10 mM HEPES.

### Traction Force Microscopy

We measured the mechanical outputs of colonies using traction force microscopy (TFM), a technique for obtaining the stress field exerted by adherent cells on their environment (Sabass et al., 2008). In a TFM experiment, cells adhere to a thick, flexible substrate with embedded fiducial markers. For our experiments, we used polyacrylamide (PAA) gels for their ease of preparation and versatility in stiffness, prepared with 0.11 µm fluorescent microspheres as markers. While cells are attached, they contract inward and deform the gel. By imaging the beads underneath cells while they are attached and subsequent to removal via SDS, we can obtain the displacement of the gel due to cell traction stresses. Based on the substrate displacement, we use Fourier Transform Traction Cytometry (Sabass et al., 2008, Butler et al., 2002) to ascertain both the magnitude and location of traction stresses, as well as derive bulk quantities such as the total mechanical output of a colony. TFM thus provides an effective means for measuring the mechanical characteristics of colonies and tissues, which makes it an attractive technique for experimentally testing predictions made by mechanical models of cells and colonies. As TFM is compatible with other imaging techniques, it also allows us to directly compare morphological changes and mechanical outputs.

### Annular Analysis for TFM

To obtain strain energy as a function of distance from the colony periphery, we adopted a similar approach to that used in (Mertz et al., 2013). Starting from a mask of the colony outline, we eroded each outline by a distance × defined in pixels; for our purposes, we used an erosion factor of 15, which corresponded to 2.42 µm. We generated a new mask consisting of the eroded region, and used this as the area over which we computed the strain energy, which is given by 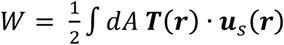, where ***T*(*r*)** is the traction stress at point **r** and ***u***_*s*_(***r***) is the substrate surface displacement at the same point. Because the TFM routine has a finite resolution and our settings generate traction footprints with radii of 5 µm or more, we commonly observe stresses exterior to the colony. We therefore included negative Δ, obtained by dilating the original outline mask, to incorporate these stresses. As Δ increases, the regions being analyzed have progressively smaller areas, so we divided strain energies by the area of each corresponding region to find the strain energy density, which decreases solely due to peripheral localization of stresses. As previously mentioned, this normalization to area meant that the strain energy density of regions close to the colony center can diverge, so we ignored any region with an area below 50 µm^2^. Finally, to facilitate comparison between different colonies, we normalized all values to Δ = 0, such that a colony with good localization to the colony periphery should have values near 1 at the edge, and 0 elsewhere.

We diverge from this process when computing the decay constant for strain energy with respect to Δ, because this quantity tends to increase in colonies with the least peripheral localization, making them unamenable to exponential fitting. We therefore used the strain energy with no normalization to area to extract these values. The strain energy profiles generated this way uniformly decrease with Δ, ensuring that they follow an exponential form. We found that the contribution from the inherent decay in stresses can still be compared to other colonies.

### Continuum mechanical model for epithelial colonies

The continuum mechanical model used here is based on our earlier work on stationary adherent cells and cohesive cell clusters (Banerjee and Marchetti, 2012; Mertz et al., 2012; Oakes et al., 2014). The model describes adherent cell clusters as a homogeneous and isotropic elastic medium, ignoring all fine-grained details regarding of individual cell mechanics and subcellular cytoskeletal architectures. The mechanical energy of the cell colony is given by 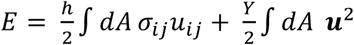. The first term in the equation characterizes the energy arising from the elastic stress tensor of the cell, which is decomposed as the sum of two parts: 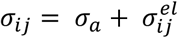, an active contractile stress and an elastic stress, respectively. In this model, *σ*^*a*^ represents the bulk contractility acting per unit area of the cell, and *σ*^*el*^ represents the elastic contribution of colony stress tensor, whose constitutive relation is governed by 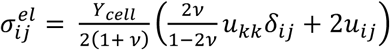. In this equation, *Y*_*cell*_ is the colony Young’s modulus and *v* is the Poisson ratio. The second term in the energy describes the energetic cost due to adhesions with the elastic substrate, where Y is an effective substrate rigidity related to the substrate stiffness and focal adhesion strength, given by 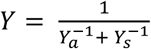 for adhesion stiffness Y_*a*_ and substrate stiffness Y_s_. This model successfully characterized the traction stress magnitude and localization in approximately circular colonies of keratinocytes, and of circular single cells (Mertz et al., 2012; Mertz et al., 2013; Oakes et al., 2014).

In the case of noncircular geometries, traction stresses are observed to localize further to regions of curvature. The model, on the other hand, smeared stresses all along the colony periphery (Oakes et al., 2014), necessitating the addition of a uniform contractile line tension, proportional to the colony perimeter, *f*_*m*_*P* This yields the energy functional: 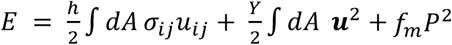, which includes the three tunable parameters of *Ycell*, *σ*_*a*_ and *f*_*m*_, with inputs of colony shape (with area A and perimeter P), colony height h, Poisson ratio *v*, and substrate rigidity Y. The resultant colony shape in steady state are determined by minimizing the total mechanical energy of the adherent colony. The traction stresses **T** can then be obtained by ***T*** = *Y****u***, where Y is the effective substrate rigidity defined above and **u** is the cell displacement field.

The values for *Y*_*cell*_, *σ*_*a*_ and *f*_*m*_,were obtained by sweeping a range of values around the values that were previously reported for single cells (Oakes et al., 2014). The resulting traction stresses were compared to experimental results on the basis of strain energy, maximum stress, and the visual extent of localization to curved regions for each geometry. The reported values were the closest to the experimental results in all three categories.

### Active Adherent Vertex Model (AAVM) for dynamic epithelial colonies

We model the cell colony as a two-dimensional monolayer. Each cell is represented by a two-dimensional polygon with mechanical energy given by 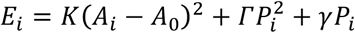 (Farhadifar et al., 2007), where *A*_*i*_ is the cell area, *A*_0_is the preferred cell area, and *P*_*i*_the cell perimeter. The first term represents the monolayers volume incompressibility and resistance to changes in height, resulting in an area elasticity; the second term represents active contractility in the actomyosin cortex; the third term represents interfacial energy due to cell-cell adhesion and cortical tension.

The mechanical energy a cell can be rearranged as: *E*_*i*_ = *K*(*A*_*i*_ – *A*_*0*_)^2^+ *Γ* (*P*_*i*_ – *P*_*0*_)^2^ (Bi et al., 2015), where 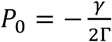 is the preferred perimeter of the cells. This gives rise to a dimensionless preferred shape index, 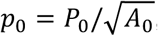, that describes the shape anisotropy of the cell. The lowest possible shape index is that of a circle, 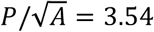. At lower values of the shape index the cells are under stress and energy is minimized by a tissue composed of isotropic, hexagonal cells. For a cell rearrangement to occur, cells must be deformed and energy temporarily increased, resulting in an energy barrier to rearrangements. As the preferred shape index increases, the energy barrier lowers until a critical shape index, 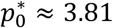, where cells can rearrange with no energy cost (Bi et al., 2015). Thus, for low preferred shape index, *p*_0_ < 3.81 (Bi et al., 2015), cortical tension dominates, the cells are hexagonal, and the tissue acts like a solid. At high preferred shape index, *p*_0_ > 3.81, cell-cell adhesion dominates, and the tissue acts like a fluid with cells able to rearrange without energy cost.

The substrate is modelled by a dynamic triangular mesh of linear springs, with focal adhesion complexes, modelled as stiff springs connecting cell vertices to substrate vertices, with adhesion energy 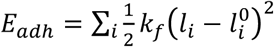, where *k*_*f*_ is the adhesion stiffness, *l*_*i*_ is the length of adhesion *i* and 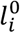 is the rest length. The cell-substrate adhesions bind and unbind stochastically. During a binding event, the adhesion springs connect the cell vertices to the nearest node of the substrate mesh, and the rest length of the adhesion bond is set to its initial length upon binding. We mimic the effects of the micropattern by disabling binding outside a chosen region of the substrate representing the micropattern geometry. Cells in the bulk move by self-propulsion with speed *v*_0_ along their polarity vector. Cells on the boundary of the colony are able to protrude their external edges, such that cell vertices are pushed forwards with a force 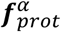 before binding to the substrate. In addition, external edges have an increased line tension, *γ*_*ext*_, to describe the preference for cell-cell adhesion over free edges.

The equation of motion for each cell vertex *α* is given by:

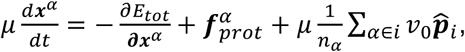

where *μ* is the viscous drag coefficient, 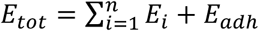 is the total mechanical energy of the cells and the cell-substrate adhesions, 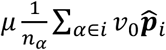 is the average self-propulsion velocity from the neighbors of vertex *α*, and 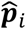 is the polarity vector of cell *i* which fluctuates over time (see Supplemental Material). The polarity vector for cells protruding out of the micropattern are reversed at a rate 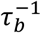.

The equation of motion for each substrate vertex *α* is given by:

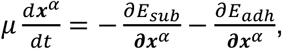

where 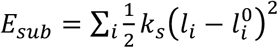 is the total strain energy in the substrate, *k*_*s*_ is the substrate spring stiffness, *l*_*i*_ is the length of the edge and *l*_*i*_ is its initial length. From displacements in the substrate we can calculate traction forces (see Supplemental Material).

At each time step, we apply the equation of motion to evolve the cell and substrate vertices and update adhesions. We allow cell-cell neighbor exchanges to occur once intercellular junctions shrink beyond a critical length (see Supplemental Material) which would reduce the total mechanical energy of the system.

### Code Availability

The computer code for Active Adherent Vertex Model (AAVM) is available at: https://github.com/BanerjeeLab/AAVM

## ACKNOWLEDGEMENTS

The authors acknowledge useful discussions with M. Cristina Marchetti. SB and MFS acknowledge support from EPSRC funded PhD studentship for MFS, UCL Global Engagement Fund, and Strategic Fellowship from the UCL Institute for the Physics of Living Systems. MLG acknowledges support from ARO MURI grant W911NF1410403 and NIH GM085087.

## Abbreviations Used

TFM: Traction Force Microscopy
ECM: Extracellular Matrix
MDCK: Madin-Darby canine kidney
PAA: Polyacrylamide
ZO-1/2 dKD: Zonula Occludens-1/2 double knockdown

